# Distinct spatial distribution of potentiated dendritic spines in encoding- and recall-activated hippocampal neurons

**DOI:** 10.1101/2025.11.19.689242

**Authors:** Francesco Gobbo, Ajesh Jacob, Bruno Pinto, Marco Mainardi, Laura Cancedda, Antonino Cattaneo

## Abstract

Experimental advancements in neuroscience have identified cellular engrams — ensembles of neurons whose activation is necessary and sufficient for memory retrieval. Synaptic plasticity, including long-term potentiation, is key to memory encoding and recall, but the relationship between learning-induced dendritic spine potentiation and neuron-wide activation remains unclear. Here, we employ a post-synaptic translation-dependent potentiation reporter (SA-PSDΔVenus) and a neuronal activation reporter (ESARE-dTurquoise) to determine their spatiotemporal correlation in the mouse hippocampal CA1 following contextual fear conditioning (CFC). Potentiated spines were enriched in activated neurons, with distribution varying across CA1 layers at different phases of memory: potentiated spines were more frequent in activated neurons in *stratum oriens* and *stratum lacunosum moleculare* after CFC (encoding), while recall-activated neurons showed a larger number of potentiated spines in the *stratum radiatum*. These findings demonstrate that the relative weight and spatial distribution of potentiated synaptic inputs to hippocampal CA1 pyramidal neurons changes between the encoding and retrieval phases of memory.

## Introduction

The search for the physical substrate of memory has been a long-standing focus of extensive research (Josselyn et al., 2017). In recent decades, the development of novel genetic tools for identifying and manipulating neurons activated during learning has provided new insights into the formation, storage and retrieval of memories (Poo et al., 2016). Immediate early genes (IEGs), including *c-fos*, *Arc* and *Zif268,* are a class of genes expressed rapidly and transiently by neuronal activation (Guzowski et al., 2001; Minatohara et al., 2016). Extensive work has demonstrated the role of IEG in memory formation and consolidation (Matynia et al., 2002; Ye et al., 2017). More recently, several groups have employed the IEG promoters to express reporter proteins or opto- or chemogenetic actuators in neurons activated during learning (Reijmers et al., 2007; Garner et al., 2012; Liu et al., 2012; Denny et al., 2014; Ohkawa et al., 2015; Vetere et al., 2017). Artificial reactivation of these neurons typically elicits a measurable behavioral response consistent with memory reactivation by natural retrieval cues, such as freezing in contextual fear conditioning (CFC) (Liu et al., 2012; Roy et al., 2022). Conversely, inhibiting these neurons impairs the expression of behavioral response elicited by natural retrieval cues (Denny et al., 2014; Tanaka et al., 2014). Using the IEG-promoter-based tools, neurons activated during learning have been identified in several brain regions, including in the hippocampal formation, the amygdala, and various cortical areas (Tonegawa et al., 2015; Vetere et al., 2017).

Bidirectional modifications of synaptic strength, collectively termed synaptic plasticity, have long been considered the neural correlate of learning (Bliss and Collingridge, 1993; Andersen et al., 2007; Takeuchi et al., 2014). An increase in synaptic response strength has been reported in the hippocampus (Whitlock et al., 2006; Pavlowsky et al., 2017), and amygdala (Rogan et al., 1997) following learning. Pharmacological treatments (Morris et al., 1986) and genetic manipulations (Tsien et al., 1996; Plath et al., 2006) that disrupt synaptic plasticity also impair adaptive behaviors in response to natural retrieval cues, although whether these interventions interfere with memory acquisition (or learning), or recall is unclear. Inducing long-term depression (LTD) and long-term potentiation (LTP) in the neuronal network modified during learning impairs and reactivates memory respectively, thus supporting a causal link between synaptic plasticity and memory (Nabavi et al., 2014).

The relationship between synaptic plasticity and the ensemble(s) of neurons activated by learning is still unclear. Synaptic plasticity, detected as an increase in spine density and AMPA/NMDA receptor current ratios, has been observed in neurons activated during learning (Ryan et al., 2015; Kitamura et al., 2017; Choi et al., 2018, 2021). Pharmacological or optogenetic inhibition of synaptic plasticity in these activated neurons impairs their reactivation and the behavioral expression of the fear memory during recall (Ryan et al., 2015; Abdou et al., 2018), extending prior results (Schafe et al., 1999; Morris et al., 2006). Their reactivation probability also changes over time and is correlated with changes in their spine density (Kitamura et al., 2017; Tonegawa et al., 2018). For instance, the spine density of hippocampal neurons activated during CFC decreases with time, and the neurons are reactivated only during recall at a recent, but not at a remote, time point (Kitamura et al., 2017).

Theoretical models and experimental studies suggest that individual memories maintain separated synaptic representations even when they largely overlap at the cellular level (Kastellakis et al., 2016; Abdou et al., 2018). Of the approximately 30,000 excitatory synapses on a single neuron (Megías et al., 2001), which ensemble of synapses is activated by learning and could represent an element of a memory trace? (Poo et al., 2016; Gobbo and Cattaneo, 2020). To address this fundamental question, novel genetic tools are being engineered to identify and manipulate the synaptic correlate of a memory (Makino and Malinow, 2011; Hayashi-Takagi et al., 2015; Gobbo et al., 2017; Dore et al., 2020; Gobbo and Cattaneo, 2020; Perez-Alvarez et al., 2020; Kim et al., 2023). Recently, we developed SynActive, a tool to express any protein of interest specifically at potentiated spines by employing regulatory sequences from the *Arc* mRNA and dendritic spine-targeting peptides (Gobbo et al., 2017). Notably, the potentiated-synapse specific expression of reporter proteins under SynActive regulation is local protein-synthesis dependent (Gobbo et al., 2017). Here, we use the SynActive strategy to map potentiated dendritic spines in the hippocampal CA1 region of mice subjected to CFC. The distribution of the potentiated spines on the dendritic arbor was then compared between active or inactive neurons during learning or recall of fear memory. Furthermore, we analyzed the distribution of the potentiated spines across different layers of the hippocampal CA1 area, which receives inputs from multiple brain areas. We demonstrate that potentiated spines are preferentially located onto CA1 neurons that are activated following CFC. The overlap between potentiated spines and neuronal activation varies across different hippocampal CA1 layers in a time-dependent manner following CFC, during the encoding or the recall phases. Our results indicate a positive correlation between the potentiated spines and the neurons activated during learning or recall of fear memory. The strength of the correlation varied across different layers depending on the time at which the animals were analyzed after CFC.

## Results

### SA-PSDΔVenus is expressed at potentiated spines in primary hippocampal neurons

Previous research has demonstrated that incorporating untranslated regions (UTRs) from *Arc* mRNA facilitates the expression of an encoded reporter or actuator protein at potentiated synapses following stimulations known to induce long-term potentiation, such as focal glutamate uncaging, spatial exploration, and motor learning (Hayashi-Takagi et al., 2015; Gobbo et al., 2017; Gobbo and Cattaneo, 2020). This type of construct functions as a protein-synthesis-dependent reporter for synaptic stimulations that lead to long-term potentiation of excitatory synapses (Gobbo and Cattaneo, 2020). In addition to the RNA regulatory element, a peptide tag or a synaptic protein is fused to the reporter to enhance its retention into the dendritic spine of potentiated synapses following activity-dependent local translation (Hayashi-Takagi et al., 2015; Gobbo et al., 2017) In this study, we mapped potentiated spines employing a fusion reporter combining the synaptic protein PSD95, the fluorescent protein mVenus and the hemagglutinin (HA) tag, under the translational regulation of the 5’ and 3’ UTRs of *Arc* mRNA (Figure 1A). PDZ1 and PDZ2 domains were deleted from the PSD95 construct (PSDΔ), to minimize overexpression artifacts while maintaining its localization to dendritic spines (Supplementary Figure 1A) (Arnold and Clapham, 1999). The resulting construct, SA-PSDΔVenus, enriched the reporter protein expression at dendritic spines in primary hippocampal neuronal cultures (Figure 1B). In comparison to our previously reported constructs (Gobbo et al., 2017), which employed a peptide tag to enhance protein enrichment at postsynaptic sites, PSDΔVenus displayed a greatly reduced reporter expression in both the dendritic shaft and soma (Supplementary Figure 1A).

**Figure 1.**
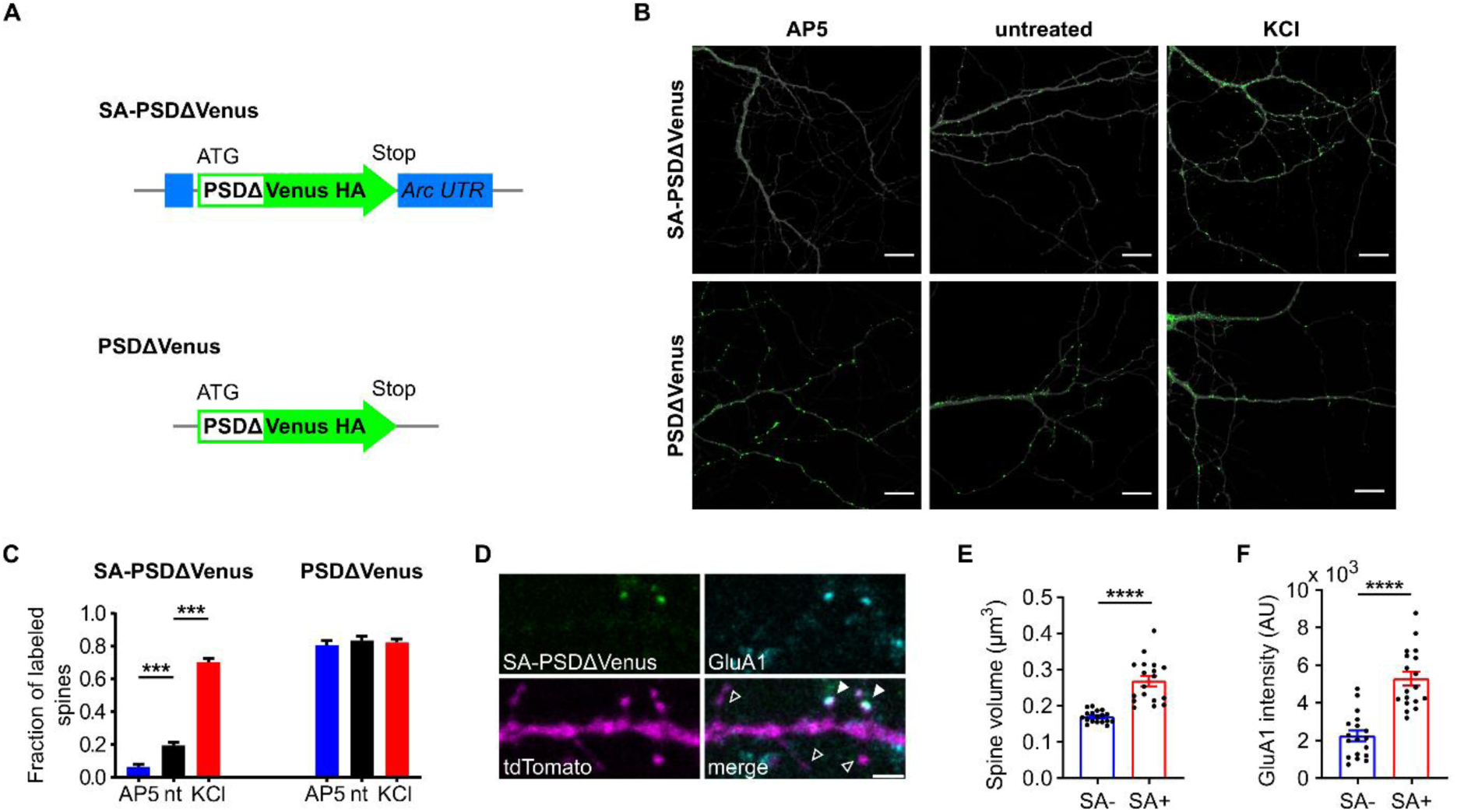
SA-PSDΔVenus is expressed at potentiated spines in an activity- and NMDA receptor-dependent manner. (A) Schema of the SA-PSDΔvenus construct and the control PSDΔVenus construct. (B) Primary mouse hippocampal neurons expressing tdTomato (greyscale) and SA-PSDΔVenus (green) or PSDΔVenus (green) after 24 hours D-AP5 (AP5), 90 minutes KCl (KCl) or untreated. Scale bar, 10 μm. (C) SA-PSDΔVenus expression is regulated by neuronal activity, while the control PSDΔVenus is not. ***P<0.001 two-way ANOVA, followed by Holm-Sidak’s multiple comparison test; interaction F(2,108)=90.96 P<0.001. (D) Hippocampal neurons expressing SA-PSDΔVenus (green) and tdTomato (magenta) immunolabelled for AMPAR subunit GluA1(cyan). The merged image shows all three channels. Filled arrowheads indicate SA-PSDΔVenus+ spines expressing GluA1, while empty arrowheads indicate spines without SA-PSDΔVenus nor GluA1. Scale bar, 2 μm.| (E and F) SA-PSDΔVenus+ spines (SA+) are potentiated – larger spine volume (E) and higher GluA1 intensity (F), compared to spines not expressing SA-PSDΔVenus (SA-) within the same tdTomato+ neurons. Circular markers indicate values corresponding to each neuron. ****P<0.0001 paired Student’s t-test, n=18 (SA- and SA+) neurons. Bar graphs show mean ± SEM (C, E and F).

In primary hippocampal cultures, KCl-induced neuronal depolarization significantly increased the fraction of SA-PSDΔVenus+ (*bona fide* potentiated) dendritic spines compared to untreated controls (Figures 1B and 1C, and Supplementary Figure 1B). In contrast, blocking synaptic potentiation with the NMDA receptor (NMDAR) inhibitor AP5 reduced the fraction of SA-PSDΔVenus+ dendritic spines. A control construct, devoid of *Arc* 5’ and 3’ UTRs, consistently expressed PSDΔVenus in nearly all dendritic spines, regardless of the treatment condition (Figures 1A-C). Furthermore, when synaptic potentiation was chemically induced using Glycine (Gly-cLTP, see Methods), we observed a larger spine volume and elevated levels of the AMPA receptor (AMPAR) subunit GluA1 in SA-PSDΔVenus+ spines, compared to non-labeled spines within the same neurons (Figures 1D-F). These are commonly used as functional and structural indicators for post-synaptic potentiation, albeit indirectly (Matsuzaki et al., 2004). These findings demonstrate that SA-PSDΔVenus is expressed at potentiated spines in an activity- and NMDAR-dependent manner.

### Contextual fear conditioning increases SA-PSDΔVenus+ spines in the hippocampal CA1 neurons

To examine the distribution of hippocampal dendritic spines potentiated *in vivo* during memory formation after a learning task, we expressed SA-PSDΔVenus in the CA1 hippocampal region of mice via triple-electrode *in utero* electroporation (Szczurkowska et al., 2016) and subjected them to contextual fear conditioning (CFC), a learning paradigm that induces strong neuronal activation and synaptic plasticity in the hippocampal regions (Takeuchi et al., 2014; Josselyn and Tonegawa, 2020). To control the temporal expression of SA-PSDΔVenus using doxycycline (dox) we employed the Tet-ON system (Zhu et al., 2007): (1) SA-PSDΔVenus under the control of the tetracycline/doxycycline-sensitive tetracycline-responsive element (TRE) promoter; and (2) the constitutively expressed reverse tetracycline transactivator (rtTA) along with a red fluorescent protein mCherry (Figure 2A). Electroporated animals received intraperitoneal injections of dox twice (the evening before and the morning of the CFC) to open the time window for labeling dendritic spines potentiated following CFC (Figure 2B). Mice subjected to CFC in context A were subsequently perfused 90 minutes (CFC group) or 24 hours (CFC24 group) after CFC. The endogenous fluorescent signals from mVenus and mCherry were comparatively weaker in hippocampal slices than in primary neuron cultures. To enhance the signal, we performed immunostaining using anti-HA and anti-mCherry antibodies to identify SA-PSDΔVenus+ spines and all spines, respectively (Supplementary Figure 2A). A control group of mice that remained in their home cages (HC) underwent the same procedures - electroporation, doxycycline injection, perfusion, and immunostaining - in parallel to CFC animals (Figure 2B).

**Figure 2.**
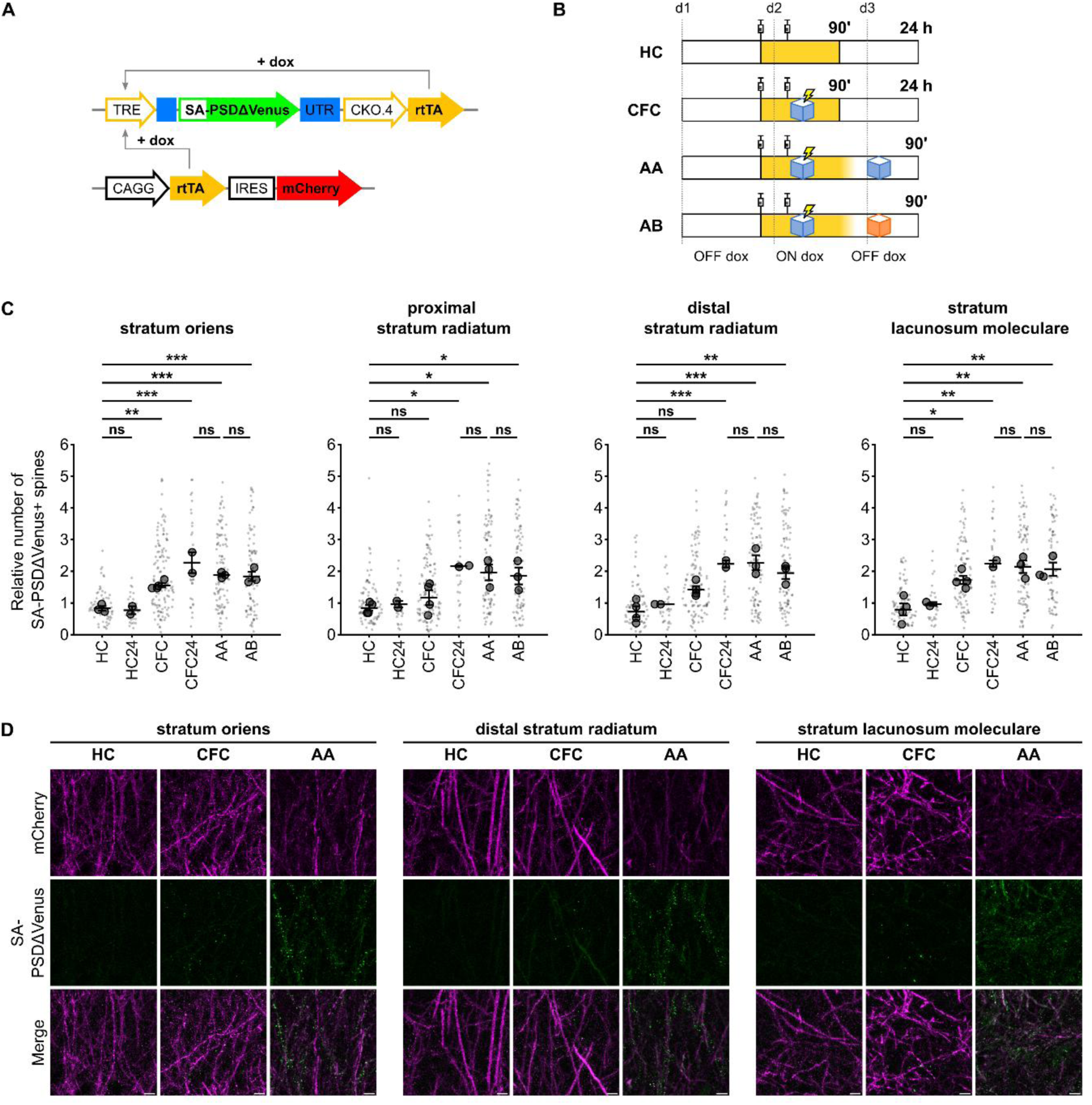
Contextual fear conditioning increases the number of SA-PSDΔVenus+ spines in the CA1 neurons. (A) Schematic of the constructs expressed in the hippocampal CA1 neurons. SA-PSDΔVenus is transcribed from tetracycline/doxycycline (dox)-sensitive TRE promoter by rtTA transcription factor (TET ON system). (B) Schematic of the four experimental groups. In yellow the time window when doxycycline is present after the injections is depicted. (C) Relative number of SA-PSDΔVenus spines in different hippocampal CA1 layers for mice remained in the home cage (HC), subjected to contextual fear conditioning (CFC) and exposed to conditioned context A (AA) or to an unrelated context B (AB) 24 hours after CFC. HC and CFC mice were perfused 90 minutes after the second dox injection, and HC24 and CFC24 animals were perfused 24 hours after CFC. See panel (B) and text for details. Bar graphs show mean ± SEM. Smaller and larger circular markers indicate values corresponding to each dendrite and the average per animal, respectively. Statistical tests were performed with N as the number of animals. *P<0.05, **P<0.01, ***P<0.001 and ns non-significant; one-way ANOVA, followed by Tukey’s multiple comparison test; SO: F(5,11)=20.69 P<0.0001, pSR: F(5,12)=6.900 P=0.0030, dSR: F(5,12)=14.53 P<0.0001, SLM: S(5,12)=12.12 P=0.0002. N=4 (HC, CFC), N=3 (AA, AB) or N=2 (HC24, CFC24) animals. (D) Representative images of *stratum oriens, stratum radiatum* and *stratum lacunosum moleculare* for the HC, CFC, and AA groups. mCherry (magenta), SA-PSDΔVenus (green) and merge are shown for each group. Scale bar, 5 μm.

Hippocampal CA1 neurons receive innervation from multiple regions of the brain (Kajiwara et al., 2008; Basu and Siegelbaum, 2015). The most distal part of the dendritic arbor (*stratum lacunosum moleculare*, SLM) receives inputs from the medial and lateral entorhinal cortex, while CA3 neurons from both hemispheres project to the dendrites in the *stratum radiatum* (SR) and *stratum oriens* (SO), which also receive sparse connections from contralateral CA1 neurons (Supplementary Figures 2B and 2C). Due to the larger area occupied by the SR, it was subdivided into two parts along the radial CA1 axis: the proximal SR (pSR), closer to the pyramidal layer, containing the pyramidal neuron somata, and the distal SR (dSR), between pSR and SLM (Supplementary Figure 2B). We quantified the fraction of SA-PSDΔVenus+ spines on the dendritic segments of hippocampal CA1 neurons across all four layers.

CFC mice exhibited a significant increase in the fraction of SA-PSDΔVenus+ spines compared to HC mice across all layers (Figures 2C and 2D, and Supplementary Figure 3). This increase was evident from 90 minutes following CFC in SO and SLM. In contrast, the increase in the pSR was less pronounced at 90 minutes but became significant after 24 hours (Figure 2C). In HC mice, which received the same two doses of dox, the fraction of SA-PSDΔVenus+ spines identified at 24 hours remained comparable to that observed at 90 minutes (Figure 2C and Supplementary Figure 3). This suggests that the time-dependent increase in SA-PSDΔVenus expression in the SR of CFC mice is unlikely to be due to background expression resulting from the extended availability of dox. Furthermore, when animals were re-exposed to the same context A (AA) or a different context (AB) 24 hours after CFC, we did not observe an additional increase in the fraction of SA-PSDΔVenus+ spines compared to CFC24 (Figures 2B-D and Supplementary Figure 3). This result indicates that SA-PSDΔVenus expression is not further induced by re-exposure to the same or a different context, consistently with the bioavailability of doxycycline, which is cleared from the brain within 24 hours after administration (Lucchetti et al., 2019).

### SA-PSDΔVenus+ potentiated spines are enriched in ESARE-dTurquoise+ active neurons in fear-conditioned mice

To investigate the distribution of potentiated synapses in activated neurons, a third DNA construct, expressing the cyan fluorescent protein mTurquoise2 under the control of the enhanced synaptic activity-responsive element (ESARE) was also co-electroporated together with the other two described above (Figure 3A). ESARE is an engineered version of the *Arc* promoter, designed to recapitulate neuronal activity-dependent gene expression from endogenous immediate-early gene (IEG) promoters (Kawashima et al., 2013). An N-terminal FLAG tag was included to amplify the mTurquoise2 signal via immunostaining. Additionally, a destabilization tag (Li et al., 1998) was fused to the C-terminal end of mTurquoise2 to reduce the protein’s half-life (dTurquoise), a strategy previously employed to align the temporal dynamics of activity-dependent protein expression with those of endogenous IEGs (Wang et al., 2006; Eguchi and Yamaguchi, 2009; Attardo et al., 2018). Consistent with these previous reports, we observed a large overlap between the expression of dTurquoise activity reporter and an endogenous IEG protein, c-fos (Figure 3B). 90 minutes after CFC, more than 80% of the hippocampal CA1 neurons immunoreactive for c-fos were also positive for ESARE-dTurquoise (Figure 3C). This implies that dTurquoise expression is a reliable proxy for neuronal activation.

**Figure 3.**
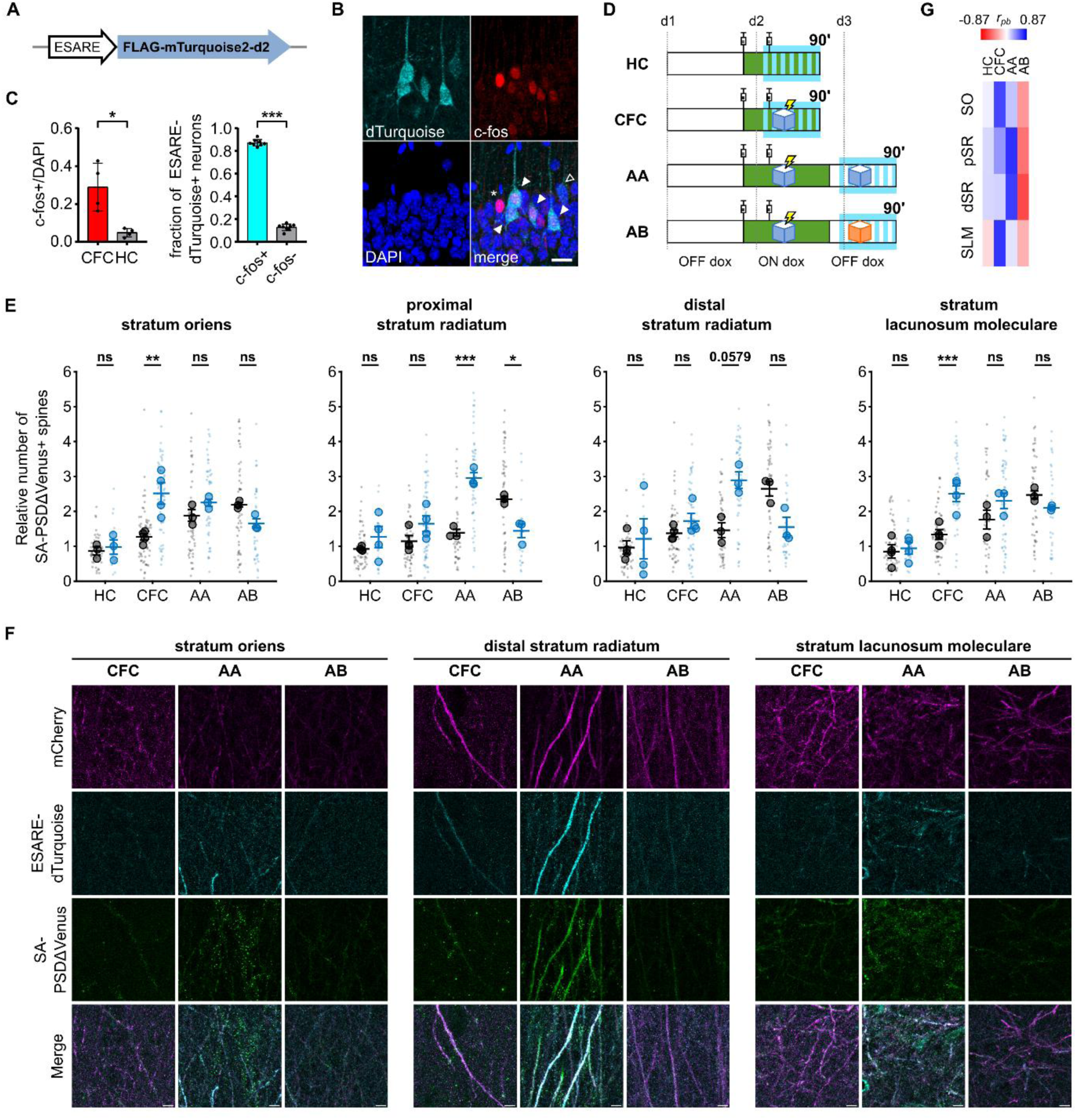
Dendritic spine potentiation overlaps with neuronal activation in the hippocampal CA1 following CFC. (A) Schematic of the construct expressed by neuronal activation. (B) ESARE-dTurquoise is expressed in c-fos+ neurons. Panels show FLAG staining (ESARE-dTurquoise), endogenous c-fos staining and DAPI channels, scale bar 20 μm. Filled arrowheads are ESARE-dTurquoise+/c-fos+ cells, an empty arrowhead shows a faint ESARE-dTurquoise+ cell without c-fos staining and an asterisk shows a c-fos+ cell without ESARE-dTurquoise expression. (C) Fraction of c-fos+ cells of all DAPI nuclei in the CFC and HC animals (left). *P=0.0297 unpaired Student’s t-test, Welch’s correction for unequal variance (t=3.757 df=3.183), N=4 animals. Fraction of ESARE-dTurquoise cells expressing c-fos in CFC animals (right). ***P<0.001 paired Student’s t-test, n=8 slices from N=4 animals. (D) Schematic of the four experimental groups. Green depicts the time window when doxycycline is present from the injections and SA-PSDΔVenus is expressed at potentiated spines. Cyan depicts the time window of ESARE-dTurquoise expression on neurons activated. (E) Relative number of SA-PSDΔVenus+ spines in ESARE-dTurquoise-(black) or ESARE-dTurquoise+ (blue) neurons in different layers as in Figure 2. Smaller and larger circular markers indicate values corresponding to each dendrite and the average per animal, respectively. Statistical tests were performed with N as the number of animals. *P<0.05, **P<0.01, ***P<0.001 and ns non-significant, two-way ANOVA, followed by Bonferroni’s multiple comparison test; SO: interaction F(3,9)=8.217 P=0.0060, pSR: interaction F(3,10)=17.09 P=0.0003, dSR: interaction F(3,10)=4.568 P=0.0291 and SLM: interaction F(3,10)=9.370 P=0.0030. N=4 (HC, CFC) or N=3 (AA, AB) animals. (F) Representative images of *stratum oriens, stratum radiatum* and *stratum lacunosum moleculare* for the CFC, AA and AB groups. mCherry (magenta), ESARE-dTurquoise (cyan), SA-PSDΔVenus (green) and merge are shown for each group. Scale bar, 5 μm. (G) Point biserial correlation (*rpb*) between the number of SA-PSDΔVenus+ spines and neuronal activation status (ESARE-dTurquoise+) in different layers for the four groups. (See also Supplementary Table 1). Bar graphs show mean ± SEM (C and E).

Next, we analyzed the distribution of SA-PSDΔVenus+ spines in hippocampal CA1 neurons that had been active (mCherry+/ESARE-dTurquoise+) or inactive (mCherry+/ ESARE-dTurquoise-) 90 minutes following CFC (Figure 3D and Supplementary Figure 4). In the SO and SLM layers, we observed a higher fraction of SA-PSDΔVenus+ spines in ESARE-dTurquoise+ neurons compared to ESARE-dTurquoise-neurons, indicating a correlation between neuronal activation and synaptic potentiation in the active neurons (Figures 3E and 3F). In control mice that remained in the home cage, the fraction of SA-PSDΔVenus+ spines was comparable between the active and inactive neurons, and the number of ESARE-dTurquoise+ neurons was quite low (Figures 3D and 3E).

We then re-exposed a group of CFC animals to the conditioned context A 24 hours after CFC (AA group). In this group, ESARE-dTurquoise is expressed by context re-exposure; conversely, SA-PSDΔVenus identifies the spines potentiated during CFC (Figure 3D). Context re-exposure does not induce SA-PSDΔVenus expression (Figures 2B-D and Supplementary Figure 3), consistent with doxycycline being cleared from the brain within 24 hours after administration (Lucchetti et al., 2019). In the pSR and dSR layers, the fraction of SA-PSDΔVenus+ spines was higher in ESARE-dTurquoise+ neurons compared to ESARE-dTurquoise-neurons (Figures 3E and 3F). This suggests that neurons harbouring a higher number of potentiated spines during the encoding phase are more likely to be reactivated during recall of that specific memory. In animals exposed to a different, non-conditioned context B 24 hours after CFC (AB group), we observed an opposite trend to AA mice (Figure 3D). Specifically, in the AB group the fraction of SA-PSDΔVenus+ spines was lower in the ESARE-dTurquoise+ neurons compared to ESARE-dTurquoise-neurons in the pSR and dSR layers (Figures 3E and 3F). This suggests that the two contexts are represented differentially at the neuronal level, and that synapses in the SR prevalently discriminate between the two, consistently with their presynaptic origin from CA3. In summary, we found a positive correlation between the neuronal activation status (ESARE-dTurquoise+) and synapse potentiation (SA-PSDΔVenus+) for the CFC and AA mice and a negative correlation for the AB mice (Figure 3G and Supplementary Table 2). The correlation in CFC mice was higher in the SO and SLM layers, while in the AA group, it was higher in the pSR and dSR layers.

## Discussion

Molecular and genetic advancement in the last two decades has led to the identification of so called cellular engrams - ensembles of neurons active at the time of learning that undergo lasting cellular changes, are necessary for memory recall and whose reactivation is sufficient for memory recall (Andersen et al., 2007; Josselyn and Tonegawa, 2020). Engram neurons are tagged via immediate early gene (IEG)-driven reporters expressed in the whole neuron. This precludes the identification of the subset of synapses linked to a particular memory, should this be the case (Andersen et al., 2007; Takeuchi et al., 2014). The relationship between synaptic and cellular memory correlates is only starting to be elucidated (Choi et al., 2018; Kim et al., 2023). To address this fundamental problem, we took advantage of the SynActive approach, which employs the combined use of untranslated sequences (UTRs) from *Arc* mRNA and synapse-targeting peptides/proteins (Gobbo et al., 2017; Gobbo and Cattaneo, 2020). Here, we report the activity-dependent expression of SynActive reporter for potentiated spines (SA-PSDΔVenus) and neuronal activity reporter (ESARE-dTurquoise) in response to an associative learning paradigm. In primary hippocampal neurons, expression of SA-PSDΔVenus was NMDAR-dependent and spatially restricted to potentiated spines (Figures 1B-F). To label the activated neurons, we adapted a previously established strategy (Li et al., 1998; Wang et al., 2006; Eguchi and Yamaguchi, 2009; Attardo et al., 2018) and expressed destabilized mTurquoise2 (dTurquoise) under the control of the enhanced synaptic activity-responsive element (ESARE) promoter (Kawashima et al., 2013). Following CFC, we observed a significant overlap between ESARE-dTurquoise expression and the endogenous neuronal activity marker c-fos (Figures 3B and 3C).

Structural and electrophysiological analysis have identified increased dendritic spine density, higher AMPAR/NMDAR current ratio, and LTP occlusion on the hippocampal neurons activated during CFC compared to inactive neurons (Takahashi et al., 2012; Ryan et al., 2015; Choi et al., 2018). This suggests preferential potentiation of spines on the neurons activated during CFC, but the individual identification of learning-induced potentiated spines *in vivo* has not been possible so far. To examine this, we labeled potentiated spines, and the neurons activated in the hippocampal CA1 region following CFC, via the expression of SA-PSDΔVenus and ESARE-dTurquoise, respectively (Figure 3D). CFC mice exhibited a significant increase in the fraction of SA-PSDΔVenus+ spines, compared to mice that remained in the home cage (HC) (Figures 2C and 2D), which, interestingly, was heterogeneous between layers. In the *stratum oriens* (SO) and *stratum lacunosum-moleculare* (SLM) layers of the CFC mice, the fraction of SA-PSDΔVenus+ spines was higher in the ESARE-dTurquoise+ neurons than in the ESARE-dTurquoise-neurons (Figure 3F), while in the proximal and distal stratum radiatum (SR), we observed a similar trend, although not statistically significant. In the SR, an increase in the fraction of SA-PSDΔVenus+ spines was detected 24 hours, but not 90 minutes, after CFC (Figure 2C). Prior research indicates that the consolidation of recently acquired memories depends on the reactivation of CA1 neurons during sharp-wave ripples recorded along the CA3-CA1 hippocampal circuit during slow-wave sleep and awake states (Sadowski et al., 2016; Wu et al., 2017; Yang et al., 2024). Based on this, we speculate that the time-dependent increase we observed may be due to SA-PSDΔVenus having labeled the spines potentiated during this consolidation phase. This is also supported by the occurrence of a second wave of LTP induction during post-learning sleep (Goto et al., 2021) and by the time-course of Arc expression in the hippocampal CA1 neurons hours after plasticity-inducing stimulation (Nakayama et al., 2015; Das et al., 2023), which has been linked to the persistence of the fear memory.

It should be pointed out that reporting “active synapses” is not the same as reporting “potentiated synapses”, undergoing translation-dependent long term potentiation after synaptic activation, the latter being currently the unique prerogative of SynActive methods (Hayashi-Takagi et al., 2015; Gobbo et al., 2017; Gobbo and Cattaneo, 2020). Regardless of this distinction, using a photoconvertible reporter, Perez-Alvarez et al identified active synapses in the hippocampus of mice running along a treadmill (Perez-Alvarez et al., 2020). A study by Choi *et. al*. using a dual-eGRASP (green fluorescent protein reconstitution across synaptic partners) system reports enhanced synaptic connectivity (increased spine number and size) between CA3 and CA1 neurons coactivated during CFC (Choi et al., 2018). Using pulse-chase labelling of surface AMPARs with membrane-impermeable dyes, Kim *et. al.* demonstrated that in CFC mice, but not in HC controls, CA1 neurons with higher neuronal activity (elevated levels of c-fos) also exhibit synaptic potentiation, as indicated by enhanced AMPAR exocytosis (Kim et al., 2023). In agreement with these reports, our study identified a positive correlation between neuronal activation and the location of synapse potentiation in CFC mice (Figures 3E and 3G). One technical advantage of the SynActive (SA-PSDΔVenus) over the dual-GRASP system is that it can label potentiated synapses in an active/inactive neuron, independent of the brain region from which it receives the input, which opens the possibility to obtain a whole neuron map of potentiated spines in a labeled neuron using sparse expression of SA-PSDΔVenus.

Memory retrieval involves the reactivation of the ensemble of neurons activated during learning (Liu et al., 2012; Denny et al., 2014; Josselyn and Tonegawa, 2020). However, whether memory retrieval reactivates the same neurons that contain the dendritic spines potentiated during learning is not known. As a first step towards this end, we labeled the spines potentiated in CA1 following CFC by SA-PSDΔVenus, and the neurons activated by memory retrieval via conditioned context re-exposure by ESARE-dTurquoise. We found that ESARE-dTurquoise+ neurons exhibited more SA-PSDΔVenus+ spines than ESARE-dTurquoise-neurons in the pSR and dSR layers (Figure 3E), suggesting neurons with increased spines potentiated during learning are more likely to be reactivated during the recall of that specific memory. Based on these differential patterns, we propose that the learning and recall of a contextual fear memory is mediated by a differential maturation of synaptic inputs to hippocampal CA1 pyramidal neurons. Our results agree with the previous studies reporting impairment in neuronal reactivation and memory recall when synaptic plasticity was disrupted in activated neurons by pharmacological treatments (Ryan et al., 2015), optogenetic stimulations (Abdou et al., 2018), or time-dependent memory consolidation (Kitamura et al., 2017). For instance, the critical role of sleep in the consolidation and maturation of memories has long been recognized, and several groups have demonstrated the role of epigenetic modifications, transcription and novel protein synthesis during sleep (Levenson and Sweatt, 2005; Lyons et al., 2023). Among these processes, replay of awake activity during sleep plays a crucial role in memory consolidation (Wilson and McNaughton, 1994; Girardeau et al., 2009; Yang et al., 2024). Neurons activated during learning exhibit higher reactivation than inactive neurons during post-learning sleep and are more likely to be reactivated during recall (Ghandour et al., 2019). Reactivation of neurons involved in learning during recall has been demonstrated with IEG labeling (Ramirez et al., 2014), electrophysiology (Ólafsdóttir et al., 2018) and calcium imaging (Cai et al., 2016; Gobbo et al., 2022). The overlap between the two ensembles is often incomplete. This could be due to drift (Ziv et al., 2013; Mau et al., 2020), circuit reorganization (Ko et al., 2025) or maturation of synaptic ensembles. It is also possible that the observed changes are, in part, due to the involvement of distinct groups of synapses. For instance, it has been demonstrated that hippocampal synapses are molecularly heterogeneous (Zhu et al., 2018) and vGlut1+ and vGlut2+ hippocampal-retrosplenial synapses contribute differentially to the formation and persistence of CFC memories (Yamawaki et al., 2019).

Notwithstanding the potential of the SynActive strategy to investigate the synaptic basis of associative memories *in vivo*, and the novelty of the *in vivo* proof of principle provided in this paper, some of the limitations of the study need to be highlighted. First, it is unclear whether the number of potentiated synapses in a particular dendritic compartment could have a preferential role in dictating the strength of memory recall (e.g. freezing levels). Second, while the available information from the literature predicts a half-life of a few hours for the destabilized reporter (Li et al., 1998; Eguchi and Yamaguchi, 2009), a more detailed evaluation of the time course for degradation and diffusion to the most distal dendrites for the reporters is needed. Third, although the spatial component of the memory is expected to be qualitatively similar in CFC animals receiving different intensities of foot shock and non-shocked animals that spent time in the conditioning context but never received any shocks, the strength of the memory can vary among the groups and correlates with the synaptic connectivity between CA3 and CA1 neurons co-activated during learning (Choi et al., 2018). How the relative number of potentiated synapses may differ between these experimental groups is unknown. Adapting the SynActive approach to a viral-based transgene delivery, which is less technically demanding than *in utero* electroporation, could facilitate the use of the SynActive approach by broader scientific community to address outstanding questions in the memory field. In any case, this new type of detailed information on the locus of synaptic plasticity following behavioral learning tasks was lacking and can be used to refine the computational models of dendritic integration and synaptic plasticity (Kastellakis et al., 2016). Furthermore, this novel approach bridges the gap between experimental methodologies with cellular and translation-dependent synaptic resolution to investigate the underlying mechanisms of memory formation and storage in both physiological and pathological contexts.

## Material and methods

### Animals

Embryonic day (E) 15.5 timed-pregnant CD1 mice (Charles River SRL, Italy) were used for the *in vivo* experiments. Time-pregnant matings were performed in the evening; the day after mating was defined as E0.5 and the day of birth was defined as P0. Mice were kept under a 12-hour dark-to-light cycle, with food and water *ad libitum*. Mice from both sexes were P26 on the day of the experiment and were assigned blindly between groups. Animal care and experimental procedures were approved with the Italian Institute of Technology licensing and the Italian Ministry of Health. Primary hippocampal neurons were prepared from P0 B6126 mice. All animal procedures were approved by the Italian Ministry of Health and Italian National Research Council (CNR).

### Plasmids

To generate PSDΔVenus we cloned PSDΔ (rat PSD95; NCBI ID NM_019621.2, nucleotides 57-248 and 993-2228) (Hayashi-Takagi et al., 2015) by joining two PCR fragments in fusion with fluorescent protein mVenus (Nagai et al., 2002) (see also Supplementary Table 1). PSDΔ was PCR amplified from FU-dio PSD95-mCherry-W (Addgene 73919). pCMV-PSDΔVenus was generated by inserting PSDΔVenus into pcDNA3.1(+) using NheI/XbaI restriction sites. pCMV-SA-PSDΔVenus was generated analogously by inserting PSDΔVenus between the 5’ and 3’ *Arc* UTRs in pcDNA3.1(+) using NheI/XhoI sites as previously described (Gobbo et al., 2017). 5’ and 3’ *Arc* UTR comprises nucleotides 1-230 and 1424–3026 of rat Arc transcript (NCBI NM_019361.2) (see also Supplementary Table 1). To generate pTRE3-SA-PSDΔVenus-HA-CK-rtTA, we first inserted the cDNA for the haemagglutinin (HA) tag in-frame at the end of the PSDΔVenus cDNA in pCMV-SA-PSDΔVenus. Then amplified by PCR the SA-PSDΔVenus-HA cDNA containing the *Arc* UTR sequences and cloned in pTRE3-SA-CK-rtTA (Gobbo et al., 2017) after removing the previous coding sequence. The resulting vector expresses SA-PSDΔVenus (PSDΔVenus-HA flanked by 5’ and 3’ *Arc* UTRs) from the third-generation tetracycline sensitive promoter (Sato et al., 2013) and the rtTA2S-M2 transactivator from the minimal CamKII(0.4) promoter. To generate pAAV-TRE3-SA-PSDΔVenus, TRE3 and SA-PSDΔVenus were cloned into pAAV-hSyn-EGFP (Addgene 50465), by replacing the sequences between the inverted terminal repeats (ITRs).

pCAGGS-TdTomato (Szczurkowska et al., 2016) expresses TdTomato from CAGGS promoter and pCAGGS-rtTA-IRES-mCherry (Gobbo et al., 2017), express tetracycline/doxycycline sensitive TetON transcription factor (rtTA) and TdTomato. pAAV-hSyn-rtTA-P2A-tdTomato was generated by cloning rtTA-P2A (P2A sequence in the 3’ end) and tdTomato between the hSyn promoter and WPRE of pAAV-hSyn-EGFP (Addgene 50465).

pESARE-dTurquoise was generated from pCAGGS backbone after removing the CAGGS-IRES-TdTomato cassette (thus leaving the polyA site) and inserting the cDNA encoding the activity-dependent promoter E-SARE (Kawashima et al., 2013) and the fusion protein FLAG-mTurquoise2-d2. The E-SARE sequence contains five copies of the SARE enhancer and was generated as described previously (Kawashima et al., 2013). To generate the FLAG-mTurquoise2-d2, the cDNA encoding mTurquoise2 was amplified from pPalmitoyl-mTurquoise2 (Addgene 36209) and inserting the cDNA for the 3xFLAG tag (*N-*DYKDHDGDYKDHDIDYKDDDDK*-C*) at the 5’ end and the cDNA encoding the destabilization sequence (*N-*HGFPPEVEEQDDGTLPMSCAQESGMDRH*-C*) from mouse ornithine decarboxylase (Li et al., 1998) at the 3’end.

### Cell culture experiments

Primary hippocampal neurons were prepared from P0 B6126 mice as previously described (Gobbo et al., 2017). Hippocampi were microdissected under sterile conditions and triturated in cold calcium-free Hank’s balanced salt solution (HBSS) (Sigma-Aldrich H6648) 100 U/ml penicillin/0.1 mg/ml streptomycin (Thermo Fisher 15070063) and digested in 0.1% trypsin (Thermo Fisher 15090046), followed by inactivation in 10% FBS (Thermo Fisher 10500064) in DMEM (Invitrogen 11880028) 100 U/ml DNase (Sigma-Aldrich D5025). Dissociated neurons were pelleted by centrifugation at 1000 rpm for 5 min, followed by resuspension in Neurobasal-A (Thermo Fisher 10888022) supplemented with 4.5 g/L D-glucose (Sigma Aldrich G7021), 10% FBS (Thermo Fisher 10500064), 2% B27 (Thermo fisher 17504044), 1% Glutamax (Thermo Fisher 35050061), 1 mM pyruvate (Sigma Aldrich S8636) and 12.5 μM glutamate (Sigma Aldrich G5889). Neurons were seeded on 24-well plates containing 13 mm glass coverslips coated with Poly-D-Lysine (PDL,) or on PDL-coated, plasma-treated Willco dishes (Willco Wells GWST-3522). The day after seeding, the medium was changed to Neurobasal-A supplemented with 2% B27, 1% Glutamax and 10 μg/ml gentamicin (neuronal growth medium). On the day in vitro (div) 2, 2.5 μM AraC was added to reduce glia proliferation. The culture medium was refreshed every 2 days.

On div 10 neurons were transfected with pCMV-SA-PSDΔVenus /pCAGGS-TdTomato or pCMV-PSDΔVenus/pCAGGS-TdTomato with calcium phosphate method. Neurons were transfected with the calcium phosphate method the day before the experiment: 10μg DNA is dissolved in 100μl 250 mM CaCl2 (Sigma Aldrich C3306), then 100μl of 2xHBS (280 mM NaCl (Sigma Aldrich S9888), 50 mM HEPES (Sigma Aldrich H4034), 1.4 mM Na2HPO4 (Sigma Aldrich S7907) pH 7.1) are added dropwise while vortexing. After 20 minutes the resulting suspension is added to the well. After 90 minutes, the medium is removed, cultures washed with 1mM MgCl2 2mM CaCl2 HBSS and 1:1 fresh:conditioned culture medium is added to the neurons. The following day neurons were treated with 10mM KCl (Sigma Aldrich P9333) for 90’ minutes or equivalent volume in saline. A third group was incubated with 100μm AP5 (Tocris 0106) overnight from the end of transfection. Neurons were then fixed in 2% PFA (Sigma Aldrich P6148) 0.5% sucrose (Sigma Aldrich S7903) in phosphate-buffered saline (PBS) for 10 minutes, then washed and maintained in PBS. Neurons were imaged using a confocal microscope (Leica TCS SP5 on DM6000, equipped with MSD module) using an oil objective HCX PL APO CS 40.0X (NA=1.25), and the pinhole was set to 1 Airy Unit (AU). The following xlaser lines/acquisition windows were used: Ar 514nm / (520/540nm) and Ar 543nm / (650/700nm) for Venus and TdTomato respectively. 1024×1024 images at digital zoom 5 were taken with 0.08μm pixel size.

For colocalization analysis of SA-PSDΔVenus and GluA1, div 3 neurons were transfected with pAAV-TRE3-SA-PSDΔVenus/pAAV-hSyn-rtTA-P2A-tdTomato using Lipofectamine 2000 (Thermo Fisher 11668019) according to the manufacturer instructions. The protocol followed for a well in a 24-well plate is as follows. The conditioned medium was collected from the well and neurons were washed with warm HBSS containing 2 mM CaCl2 and 1 mM MgCl2. Neurons were then placed back in the incubator after adding 400 µl of fresh neuronal growth medium. pAAV-TRE3-SA-PSDΔVenus (1 µg)/pAAV-hSyn-rtTA-P2A-tdTomato (1µg) plasmid mix was then diluted in Opti-MEM to 50 µl and mixed with Lipofectamine 2000 (3 µl) diluted in 50 µl Opti-MEM. After incubation at room temperature for 5 minutes, the 100 µl transfection mix was added to the neurons and incubated for 2.5 hours at 37°C. Finally, the transfection mix-containing medium was replaced with the conditioned medium (350 µl) and fresh neuronal growth medium (150 µl). Doxycycline (Sigma-Aldrich D9891, final concentration 1 μg/ml) was added to cultures the evening before the induction of Glycine-mediated chemical LTP (Gly-cLTP), between div14 −16. Gly-cLTP was induced in cultured hippocampal neurons as previously described (Lu et al., 2001). Briefly, conditioned medium was collected, and the neurons were initially incubated in extracellular solution (ECS, pH 7.4, containing 140 mM NaCl, 1.3 mM CaCl2, 5 mM KCl, 25 mM HEPES, 33 mM D-glucose, 0.5 µM tetrodotoxin (TTX, Tocris 1078), 1 µM strychnine (Sigma-Aldrich) and 50 µM picrotoxin (PTX, Tocris 1128) at 37°C for 30 min. The solution was then replaced with ECS containing 200 µM Glycine (Sigma-Aldrich) to induce LTP. After 3 min, the solution was switched back to normal ECS and neurons were incubated at 37°C for another 30 min. Finally, ECS was removed, and the conditioned medium aspirated at the beginning of the experiment was added back. The neurons remained in the 37°C incubator until fixation at 90 minutes post Gly-cLTP. After fixation, neurons were permeabilized in 0.1% Triton X-100, 2.5% bovine serum albumin (BSA) PBS for 7 min, followed by five washes with PBS and blocking in 5% BSA PBS for 1 h. Neurons were then incubated with rabbit monoclonal GluA1 antibody (Cell signaling 13185, 1:500) in 2.5% BSA PBS for 2-3 h at room temperature. After washing thrice with PBS, neurons were incubated with anti-rabbit-647 (Thermo Fisher A-31573) in 2.5% BSA PBS for 1 h at room temperature. Neurons were washed thrice in PBS and once in the water, and the glass coverslips with fixed neurons were then mounted using Vectashield antifade (Vector Laboratories) mounting media. Neurons were imaged using Zeiss LSM 800 (oil objective - Plan-Apochromat 63×/1.4 NA DIC M27) confocal microscopes and the pinhole was set to 1 Airy Unit (AU). The following laser lines were used: 640 nm, 561 nm and 488 nm for GluA1, TdTomato and mVenus, respectively. 1024×1024 images were taken with 0.099 μm pixel size.

### *In utero* electroporation and animal experiments

Hippocampal *in utero* electroporation was performed as previously described (Szczurkowska et al., 2016). Embryonic day (E) 15.5 timed-pregnant CD1 mice (Charles River SRL, Italy) were used. Time-pregnant matings were performed in the evening; the day after mating was defined as E0.5 and the day of birth was defined as P0. Embryos from time-pregnant mothers were electroporated unilaterally with pTRE3-SA-PSDΔVenus-HA-CK-rtTA / pCAGGS-rtTA-IRES-mCherry / pESARE-dTurquoise. Mice from both sexes were P26 on the day of the experiment and were assigned blindly between groups. Animals received 0.5 mg doxycycline (1 mg/30 g body weight) in saline solution intraperitoneally the evening before the experiment, and early in the morning of the experiment. After 3-4 hours animals were put in the fear chamber (a square box with metal rods on the floor). After three minutes in the chamber a 2 seconds 0.75 A shock was administered through the metal floor and maintained in the chamber for 30 seconds after the shock. The CFC group was fixed by transcardial perfusion with 4% formaldehyde in phosphate-buffered saline (PBS) 90 minutes after exiting the chamber. Due to the time necessary for animal preparation and perfusion, up to 120 minutes could pass between the end of conditioning and the fixation of the brain. In the text, this time interval will be considered as 90 minutes. Mice from the HC group were injected identically but were maintained in their home cage and were not exposed to CFC. HC animals were perfused at times matched with the CFC group. Animals from the AA and AB groups were injected and conditioned as the CFC group but were returned to their home cage after CFC until the following day. 24 hours after CFC, animals from the AA group were put in the conditioned chamber for 3 minutes with no shock, while the AB animals were put in the control chamber (a square box with a different floor and walls) for the same amount of time. Both groups were perfused after 90 minutes. HC24 and CFC24 groups were injected and treated as the HC and CFC groups, respectively, but were perfused 24 hours after CFC (or equivalent time for HC24 animals) instead of 90 minutes.

### Immunofluorescence and image acquisition

After perfusion, brains were post-fixed overnight in 4% formaldehyde in PBS, then cryoprotected in 30% sucrose PBS. Sixty μm-thick coronal sections were cut with a cryostat. After washing the slices in PBS, slices were blocked for 1h in PBS 0.3% Triton X-100 (Sigma-Aldrich T8787), 10% filtered FBS (Thermo Fisher 10500064), then incubated with primary antibodies goat anti-HA (Santa Cruz, sc-805-g) 1:100 / rabbit anti mCherry (Abcam ab167453) 1:200 / mouse anti-FLAG (Sigma-Aldrich F3165) in PBS 0.3% Triton X-100, 10% FBS overnight at 4°C shaking. After incubation, slices were washed three times for 10 minutes each with PBS 10% FBS 0.1 % Triton x-100, then incubated with 1:200 donkey anti-goat Alexa Fluor488 (Thermo Fisher A-11055)/ 1:200 donkey anti-Rabbit Alexa Fluor555 (Thermo Fisher A-31572)/ 1:200 donkey anti-mouse Alexa Fluor647 (Thermo Fisher A-31571) in PBS 0.1% Triton X-100 10% FBS for three hours at room temperature upon shaking. Slices were then incubated for ten minutes in PBS Triton X-100 0.1% with 10 μg/ml DAPI (Sigma-Aldrich D9542), then washed three times in PBS for 10 minutes, and once in deionized water before mounting them in VECTASHIELD Antifade Mounting Medium (Vector Laboratories H-1000). 1024×1024 pixel images were acquired in the hippocampal area CA1 with a confocal microscope (Leica TCS SP5 on DM6000, equipped with MSD module) using an oil objective HCX PL APO CS 40.0X (NA=1.25), and the pinhole was set to 1 Airy Unit (AU). The area was identified by morphological criteria using the DAPI staining. Optical rotation was performed so that the CA1 pyramidal layer was parallel to one of the image axes, then the sample was moved so that the desired area would lie in the centre of the field of view. Digital zoom was set to 3, yielding a final pixel size of 0.13 μm. The following excitation wavelength/acquisition windows were used for the different fluorophores: DAPI (405 nm/415-465 nm) Alexa Fluor488 (488 nm/500-550 nm), Alexa Fluor555 (561 nm/565-650 nm), Alexa Fluor647 (633 nm/645,700 nm). Sequential illumination with HeNe 633, Ar 488, DPSSL 561 and diode (Picoquant, Berlin, Germany) 405 laser lines to minimize photobleaching. To compare the overlap between ESARE-dTurquoise and endogenous c-fos expression, slices from HC or CFC groups were stained for c-fos/FLAG immunoreactivity as follows: 1 h blocking in PBS 0.3% Triton X-100 (Sigma-Aldrich T8787), 10% normal goat serum (NGS) (Sigma Aldrich NS02L), then incubated with primary antibodies mouse anti-FLAG (Sigma-Aldrich F3165) / rabbit anti-c-fos (Santa Cruz, sc-52) 1:100 in PBS 0.1% Triton X-100, 10% NGS overnight at 4°C shaking. After three washes for 10 minutes each with PBS, slices were incubated with 1:200 donkey anti-mouse Alexa Fluor488 (Thermo Fisher A-21202) / 1:200 donkey anti-rabbit Alexa Fluor647 (Thermo Fisher A-31573) in PBS 0.1% Triton X-100 for three hours at room temperature upon shaking. Slices were then incubated for ten minutes in PBS Triton X-100 0.1% with 10 μg/ml DAPI (Sigma-Aldrich D9542), washed three times in PBS for 10 minutes, and once in deionized water before mounting them in VECTASHIELD Antifade Mounting Medium (Vector Laboratories H-1000).

### Data quantification

In cell culture experiments, the number of mVenus+ spines was counted and normalized by the total number of spines, counted from the tdTomato channel. AMPAR enrichment analysis was performed as previously described (Zhang et al., 2015). Briefly, circular ROIs of a radius approximately equal to the spine head (tdTomato+) were drawn over the spines with or without the SA-PSDΔVenus signal and background subtracted integrated AMPAR intensity was calculated for each ROI. Background intensities were obtained from the spine ROIs translated in x/y to a nearby region. The volume of a spine was calculated using the formula, 4/3 *πr*3, where *r* is the radius of the corresponding spine ROI.

In animal experiments, areas from different CA1 subfields were analyzed according to standard classification (Paxinos and Franklin, 1997; Andersen et al., 2007). *Stratum oriens* (SO) is defined as the tissue layer between the outer pyramidal layer and the surface of the hippocampus containing the basal dendrites; *stratum radiatum* (SR) is the layer containing the apical dendrites while the *stratum lacunosum-moleculare* (SLM) is the most distal part of CA1, comprising the last 50-60 μm before the hippocampal sulcus separating CA1 from the dentate gyrus. Morphologically, the SLM coincides with the apical tuft of CA1 pyramidal neurons (Spruston, 2008). To account for the larger area of SR compared to SO and SLM, SR was divided into a proximal part (pSR) closer to the soma layer and a distal part (dSR) closer to the SLM. Dendrites in the four sets were selected based on the mCherry channel, and then the number of SA-PSDΔVenus-positive spines was counted. The number is expressed as the number of SA-PSDΔVenus+ spines over the whole number of spines in the dendrite, as evaluated by the filler mCherry channel, normalized on the control group. Last, after counting, the dendrite was classified as ESARE-dTurquoise+ or ESARE-dTurquoise-based on the presence or absence of anti-FLAG staining in the 647 channel. Counting was performed blind to the experimental group.

### Quantification and statistical analysis

Image analysis was performed using ImageJ (Schindelin et al., 2012). Statistical analysis was performed with OriginPro v9.0 or GraphPad Prism 8. For comparing SA-PSDΔVenus and PSDΔVenus expression in primary neuronal cultures, two-way ANOVA followed by Holm-Sidak’s multiple comparison test was performed. Paired Student’s t-test was used to compare the AMPAR enrichment in primary neuronal cultures. The number of SA-PSDΔVenus+ spines in the CA1 hippocampal neurons under different experimental conditions were analyzed using one-way ANOVA, followed by Tukey’s multiple comparison test or two-way ANOVA, followed by Bonferroni’s multiple comparison test.

Statistical details and sample sizes can also be found in each Figure legend and the minimum significance level defined was *p* < 0.05. The bar graphs on the images show mean ± SEM.

## Data availability statement

All the reagents generated in this study are available from the lead contact under an agreed materials transfer agreement. Any additional information is available from the lead contact upon request.

## Ethics statement

All animal procedures were approved by the Italian Ministry of Health and the Italian National Research Council (CNR).

## Author contributions

Conceptualization, AC; Project plan, AC and FG; Experimental investigation, FG and AJ; *in utero* electroporation and *in vivo* logistics, LC and BP; Data analysis, FG and AJ; Data interpretation and discussion, AC, FG and AJ; Overall supervision, AC and MM; Writing – original draft, AC, FG, and AJ; Writing – review & editing, AC and AJ; Critical reading of final version of the manuscript, all authors.

## Acknowledgements

We thank Alessandro Viegi for expert technical and infrastructural assistance and Francesca Chiara Latini (SNS), Mariachiara Di Caprio (SNS) and Silvia Marinelli (EBRI) for useful discussions and critical reading of the manuscript.

## Supplementary Material

**Supplementary Figure 1.**
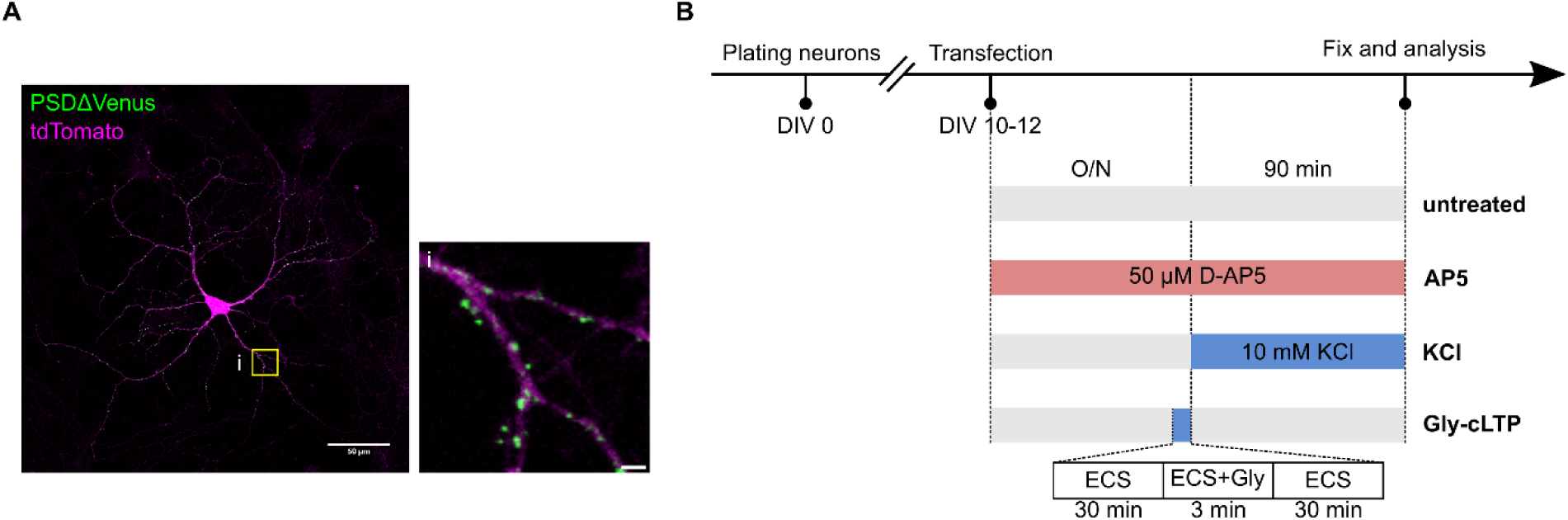
PSDΔVenus labels dendritic spines in primary hippocampal neurons. (A) Expression of PSDΔVenus in primary hippocampal neurons (green) along with tdTomato (magenta). The signal is confined to dendritic spines and absent from axons and soma. Scale bar, 50 μm; inset: 2 μm. (B) Schematic of the treatments used to modulate synaptic activity in the primary neuron cultures.

**Supplementary Figure 2.**
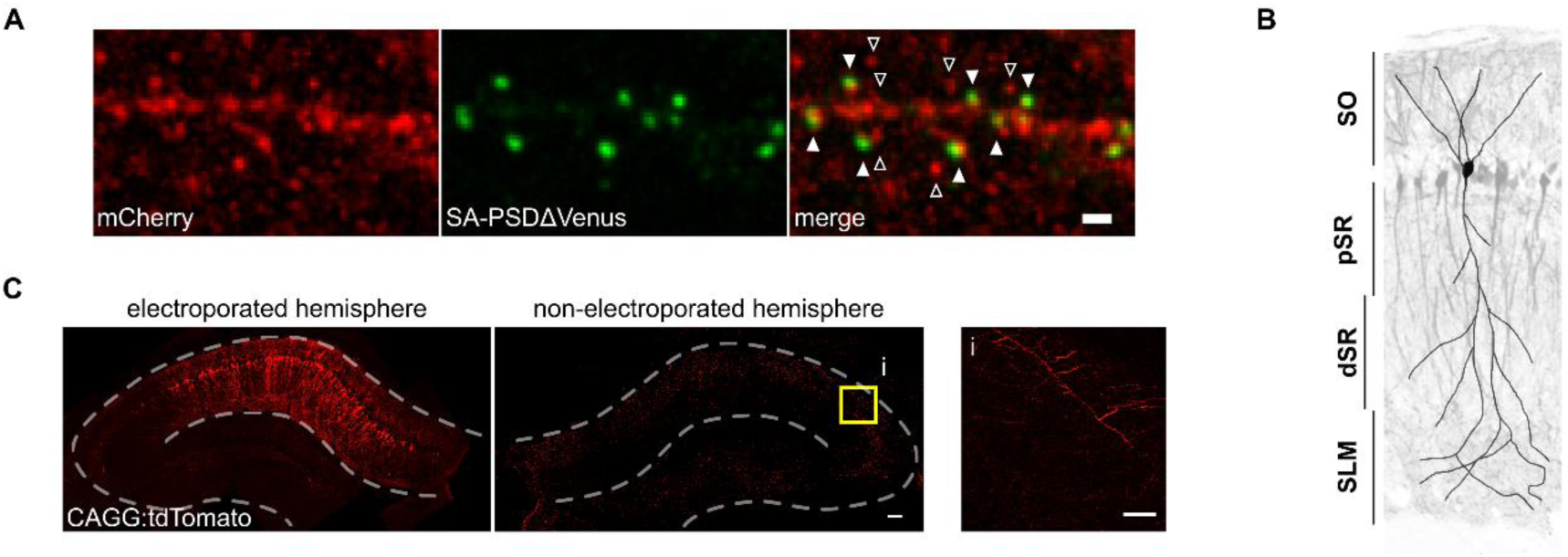
Expression of SA-PSDΔVenus and filler fluorescent proteins in the mouse hippocampal CA1 area. (A) Expression of neuron filler mCherry (red; anti-mCherry staining) and SA-PSDΔVenus (green; anti-HA staining) in the hippocampal CA1 neurons. Scale bar, 1μm. Filled arrowheads indicate SA-PSDΔVenus+ spines, while empty arrowheads are spines without SA-PSDΔVenus. (B) Schematic of the regions considered in this study in the CA1 area, superimposed on the image of a mouse electroporated unilaterally in CA1 with CAG:TdTomato. Abbreviations are: SO: *stratum oriens*, pSR and dSR: proximal and distal portion of the *stratum radiatum*, respectively; SLM: *stratum lacunosum-moleculare*. (C) Interhemispheric projection of CA1 neurons to contralateral CA1. Animals were unilaterally electroporated in CA1 with CAGG:TdTomato. Axonal projections in the non-electroporated hemisphere are prevalently confined to CA1 stratum oriens (arrowheads). An inset (i) is showed, where presynaptic boutons are evident. Scale bar, 100 μm; inset: 25 μm.

**Supplementary Figure 3.**
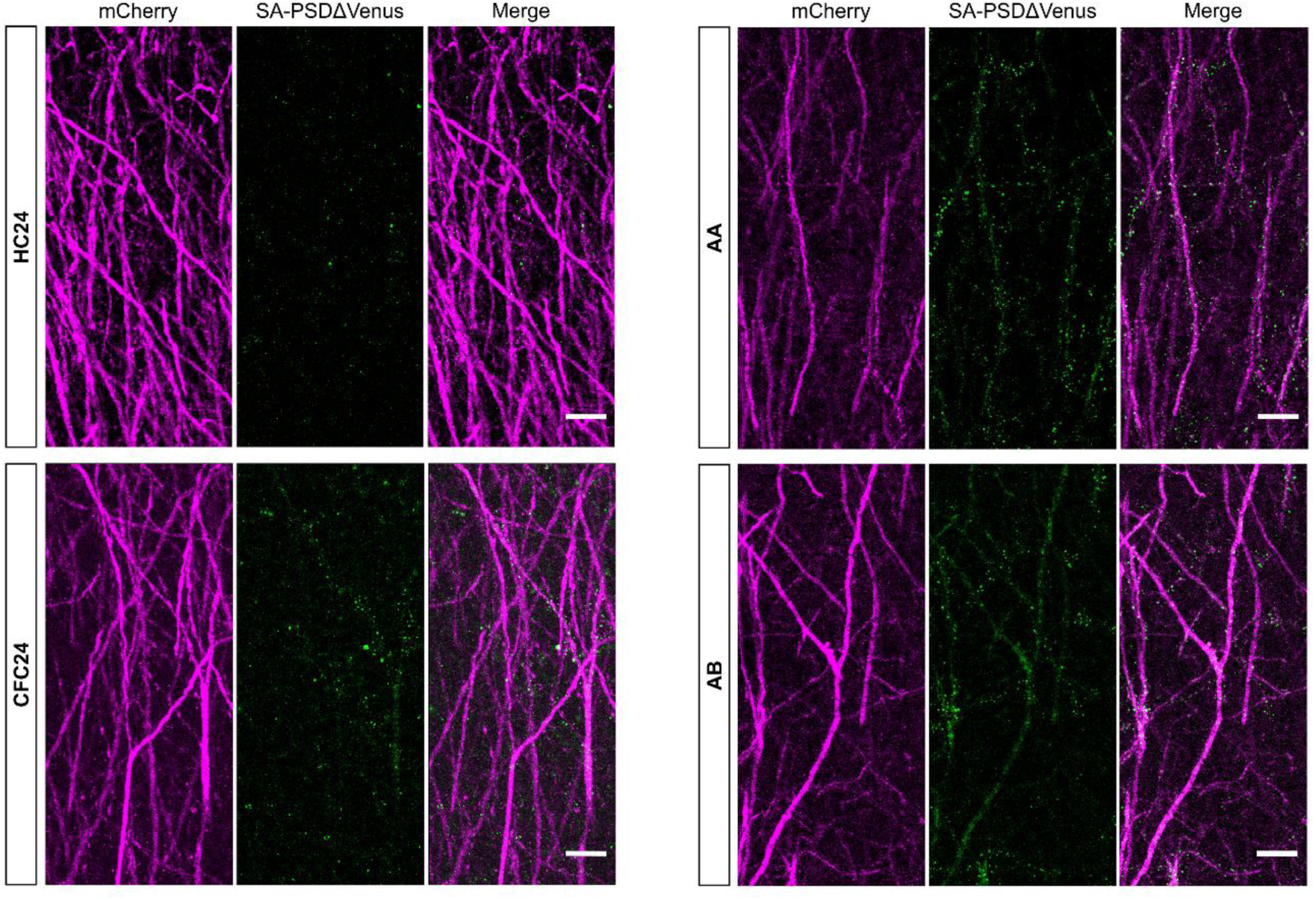
SA-PSDΔVenus expression in AA and AB mice is similar to CFC mice at 24 hours. Representative images showing dendritic SR regions for HC24, CFC24, AA and AB animals. mCherry (magenta), SA-PSDΔVenus (green) and merge are shown for each group. Scale bar, 10 μm.

**Supplementary Figure 4.**
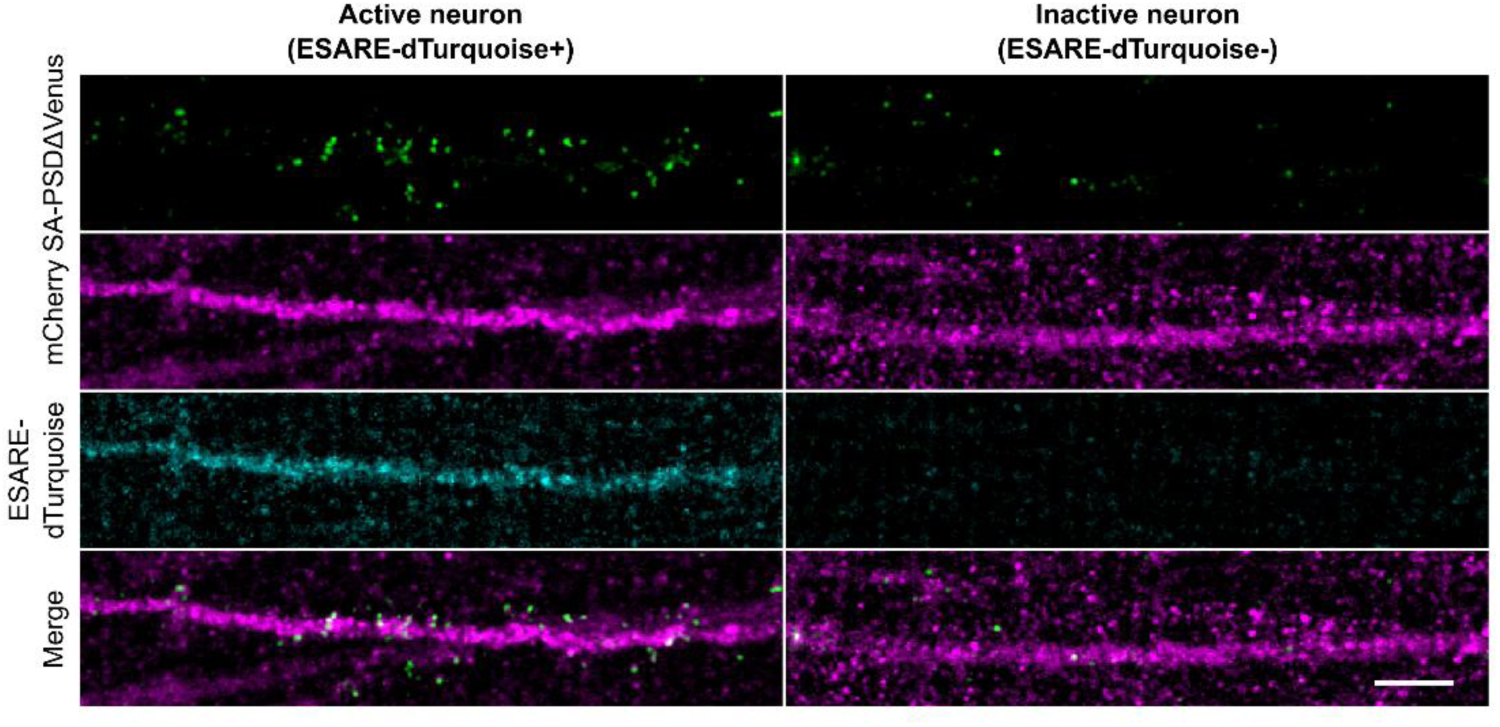
SA-PSDΔVenus+ spines in the *stratum radiatum* preferentially map onto neurons active during memory recall. Differential distribution of SA-PSDΔVenus+ spines in *stratum radiatum* dendrites of ESARE-dTurquoise+ (active) and ESARE-dTurquoise-(inactive) neurons in the AA group. SA-PSDΔVenus (green), mCherry (magenta), ESARE-dTurquoise (cyan) and merge are shown for each group. Scale bar, 5 μm.

**Supplementary Table 1.**
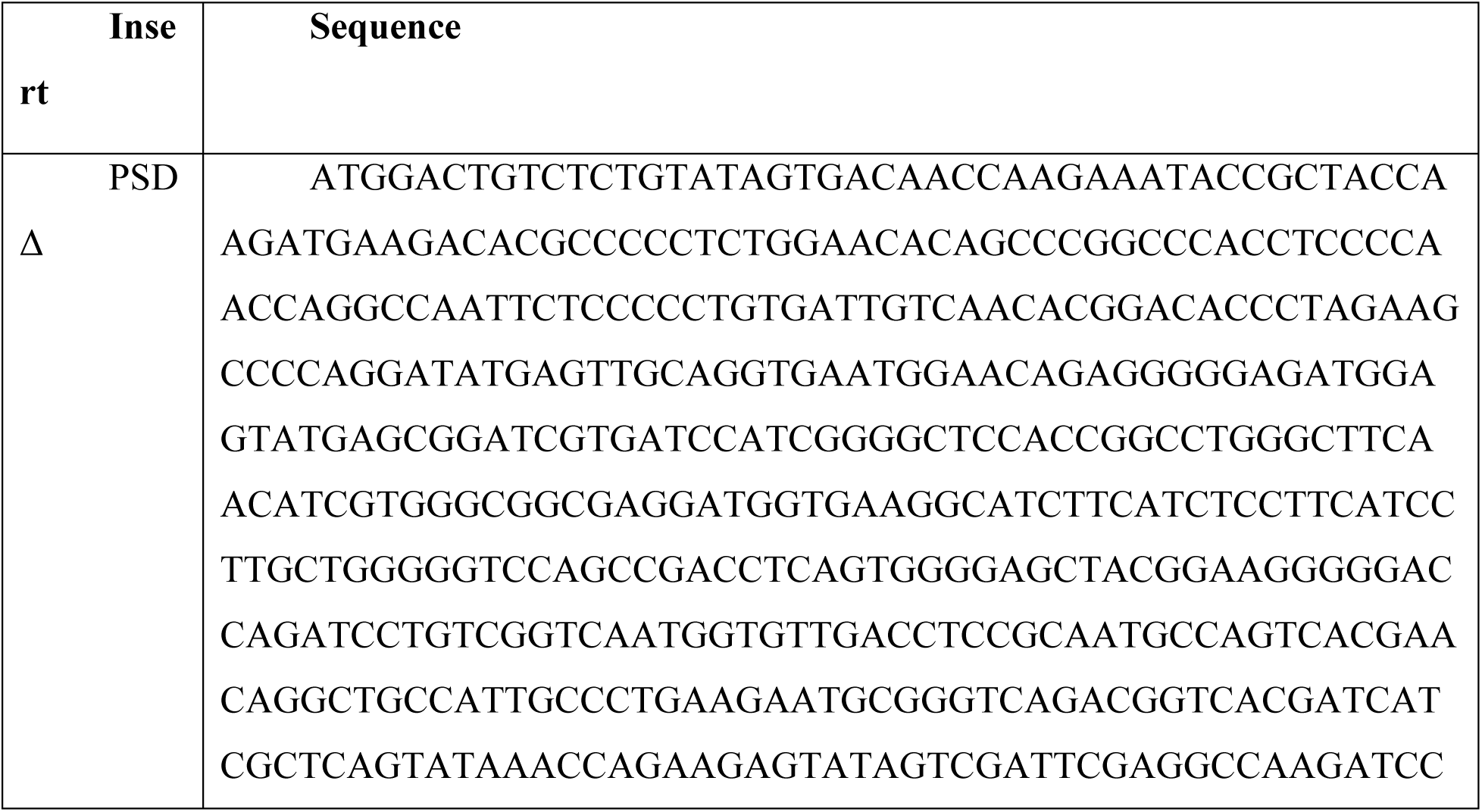
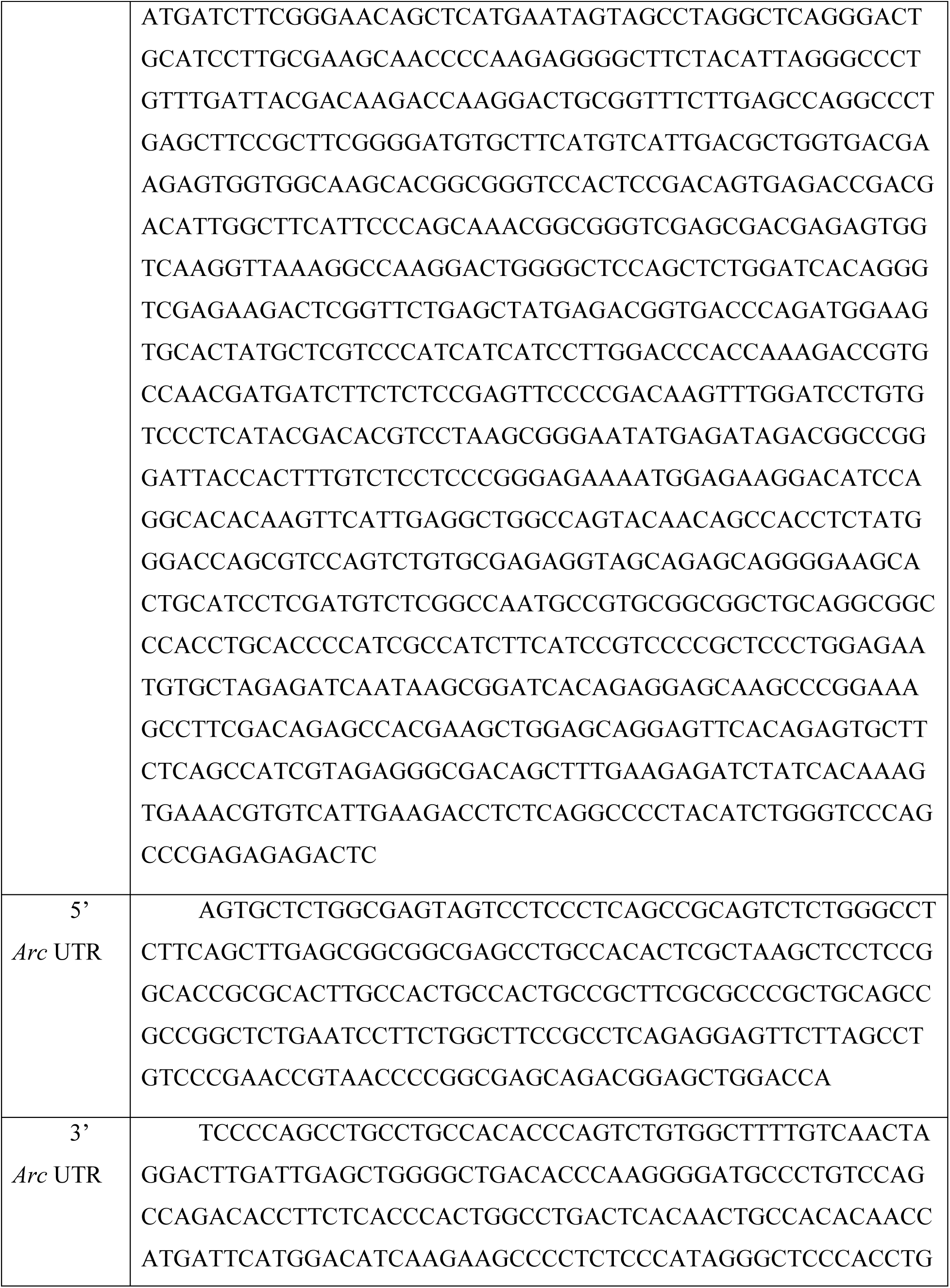
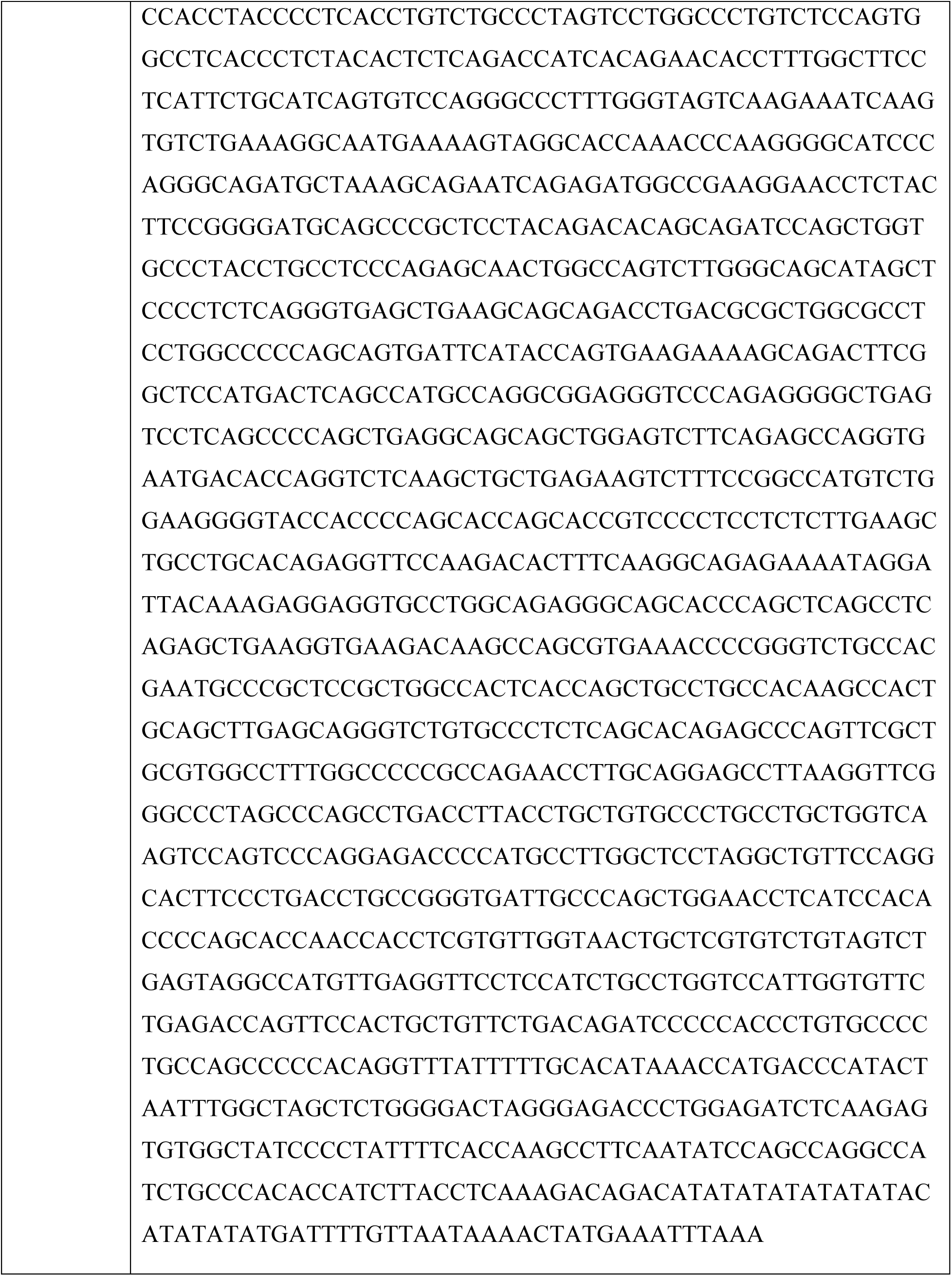
Sequences of PSDΔ, 5’ and 3’ Arc untranslated regions (UTRs).

**Supplementary Table 2.**
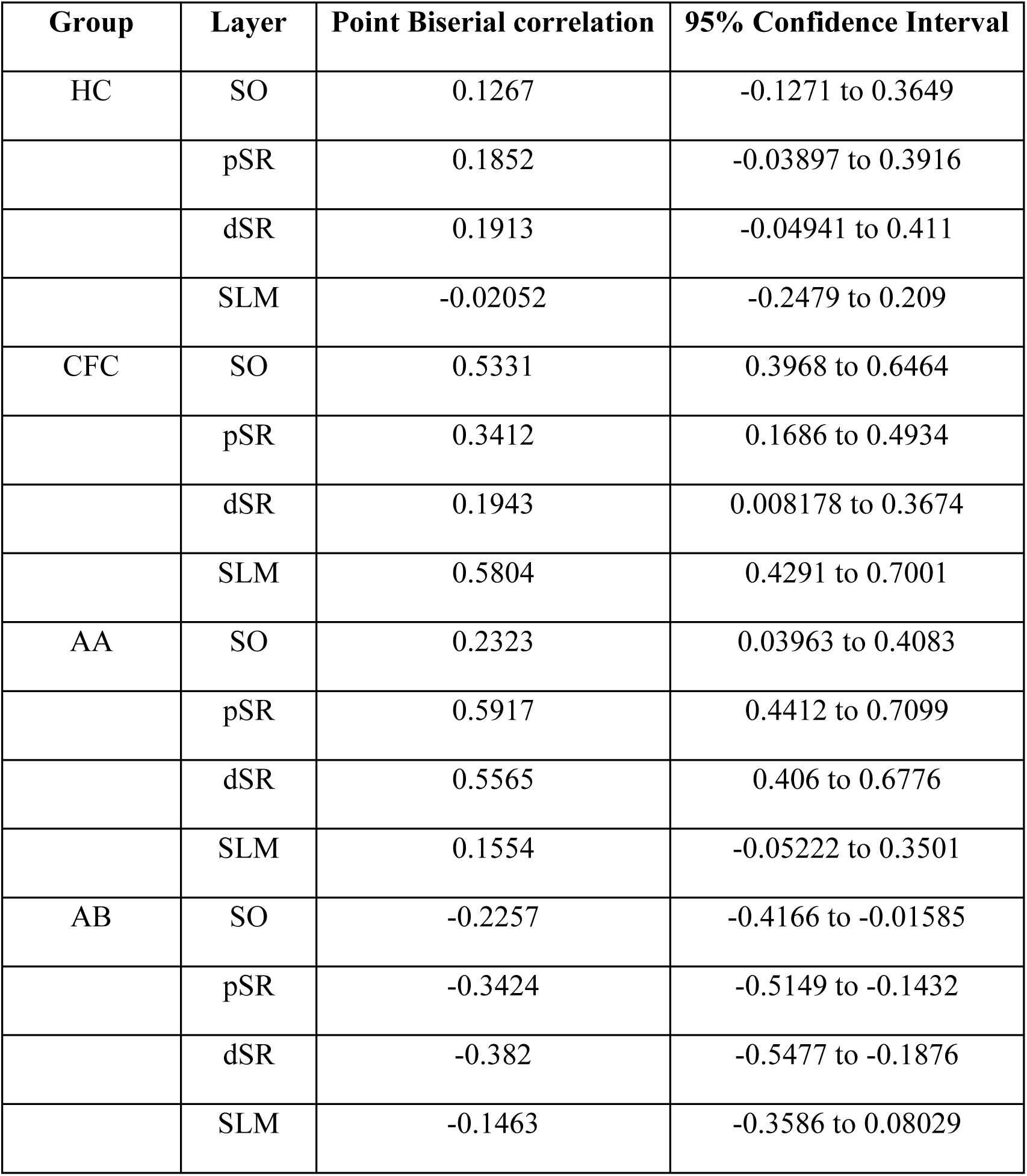
Point biserial correlation (*rpb*) between the number of SA-PSDΔVenus+ spines and neuronal activation status (ESARE-dTurquoise+) in different layers for the four experimental groups.

